# The monoplastidic bottleneck in algae and plant evolution

**DOI:** 10.1101/109975

**Authors:** Jan de Vries, Sven B. Gould

## Abstract

Plant and algae plastids evolved from the endosymbiotic integration of a cyanobacterium by a heterotrophic eukaryote. A consequence of their ancestry is that new plastids can only emerge through fission and vital to organelle and host co-evolution was the early synchronization of bacterial division with the host’s eukaryotic cell cycle. Most of the sampled algae, including multicellular macroalgae, house a single plastid per cell — or nucleus in case of coenocytic cells — and basal branching relatives of polyplastidic lineages are all monoplastidic. The latter is also true regarding embryophytes, as some non-vascular plants are monoplastidic at least at some stage of their life cycle. Here we synthesize recent advances regarding plastid division and associated proteins, including those of the peptidoglycan wall biosynthesis, across the diversity of phototrophic eukaryotes. Through the comparison of the phenotype of 131 species harbouring plastids of primary or secondary origin, we uncover that one prerequisite for an algae or plant to house multiple plastids per nucleus appears the loss of the genes MinD and MinE from the plastid genome. Housing a single plastid whose division is coupled to host cytokinesis appears a prerequisite of plastid emergence; escaping that monoplastidic bottleneck succeeded rarely and appears tied to evolving a complex morphology. Considering how little we know about the mechanisms that guarantee proper organelle (and genome) inheritance raises the peculiar possibility that a quality control checkpoint of plastid transmission remains to be explored and which is tied to understanding the monoplastidic bottleneck.

## From a free-living cyanobacterium to an inheritable plastid

The origin of plastids traces back to the endosymbiotic integration of a cyanobacterium into the cytosol and biochemistry of a heterotrophic eukaryote. We know almost nothing about the nature of the protist host and the type of cyanobacterium that engaged into this endosymbiosis is still disputed. Different analyses have uncovered different cyanobacterial phyla as the source from which the primal plastid evolved (Deschamps et al., 2008; Deusch et al., 2008; Criscuolo and Gribaldo, 2011). Based on overall sequence similarity between nuclear-encoded proteins of plants and cyanobacterial genomes, section IV or section V cyanobacteria — with the ability to fix atmospheric nitrogen — appear the plastid donor (Dagan et al., 2013), other phylogenetic analyses point to the recently discovered freshwater-dwelling *Gloeomargarita-clade* (Ponce-Toledo et al., 2017). What is commonly accepted is that the three archaeplastidal lineages, the Glaucophyta, the Rhodophyta and the Chloroplastida (Zimorski et al., 2014; Archibald, 2015), arose monophyletically (Rodríguez-Ezpeleta et al., 2005; Jackson and Reyes-Prieto, 2014; Burki, 2014). This major group emerged early during eukaryotic evolution (He et al., 2014) and most likely sometime between the ‘upper limit’ for the origin of eukaryotes ~1.9 Gya (Eme et al., 2014) and the fossilization of *Bangiomorpha* some 1.2 Gya (Butterfield, 2000). The fossil records of *Bangiomorpha* display a multicellular organism with branched filaments (Butterfield, 2000), allowing to speculate that single-celled algae, maybe comparable to extant glaucophytes, are older.

The consummated endosymbiotic integration of a prokaryote into a eukaryote is rare. While we find examples of endosymbiotic integration of prokaryotes in several clades of eukaryotes, for example the spheroid body of *Rhopalodia gibba* (Kneip et al., 2008) and *Epithemia turgida* (Nakayama et al. 2014), the chromatophore of *Paulinella chromatophora* (Nowack, 2014), or the obligate and sometimes intricate endosymbionts of many insects (Bennett and Moran, 2015; Husnik and McCutcheon 2016), we only know of two that have manifested themselves in the bigger picture of evolution: the mitochondrion and the plastid (Zimorski et al., 2014). Together they were key to the evolution of all macroscopic life. Among the numerous reasons for why such an integration is rare are the challenges of (i) fusing a prokaryote and eukaryote genome (Timmis et al., 2004), (ii) establishing retargeting of proteins now encoded in the host nucleus and translated in the cytosol as a consequence of endosymbiotic gene transfer (Soll and Schleiff, 2004; Leister, 2016; Garg and Gould, 2016), (iii) developing means of communication such as retrograde signalling (Woodson and Chory, 2008; Chandel 2015; Singh et al. 2015), and (iv) synchronizing prokaryotic (cycle-lacking) fission with the cell cycle of the eukaryotic host (Miyagishima, 2011). Organelle and host co-evolution depends on the simultaneous and successful implementation of all of these events. In some cases, however, certain steps compete with each other. These conflicts add an additional layer of complexity and in light of this manuscript this concerns in particular gene transfer from the endosymbiont to the host genome and the number of plastids per cell.

## Think big: endosymbiotic gene transfer

Endosymbiotic gene transfer (EGT) occurs not in the form of individual genes, but usually in chunks of entire genome fragments (Henze and Martin, 2001; Yuan et al., 2002; Michalovova et al., 2013). The transfer of larger fragments of DNA is thought to occur through the uncontrolled lysis of endosymbionts under certain conditions, resulting in the release of organellar DNA of which some can be randomly incorporated into the nuclear genome (Martin, 2003). Many nuclear genomes of eukaryotes carry evidence for recent transfers that are known as nuclear mitochondrial (NUMTs) and nuclear plastid (NUPTs) DNA sequences (Richly, 2004; Hazkani-Covo et al., 2010); this also sheds some light on the number of endosymbionts present, at times when the bulk of EGT occurs. If an alga carries only one endosymbiotic plastid, then its lysis results in the death of the host cell and hence no progeny. That is why EGT preferably occurs in the presence of multiple donors per cell, which is true both for today’s organelles (Smith et al., 2011; Smith, 2011) as well as to the cyanobionts that evolved into modern plastids (Barbrook et al., 2006; Fig. 1). The presence of a single organelle, whether plastid or mitochondrion, significantly slows down the rate of EGT if not bringing it to a halt altogether and freezing the organellar genomes in time (Barbrook et al., 2006; Curtis et al., 2012). The presence of multiple cyanobionts during the early stages of endosymbiosis by all means seems likely, maybe a prerequisite. At some point of their coevolution, however, it appears that the host cell and plastid needed to reach a ratio of 1:1, a ratio we find conserved in the majority of algae today and that is rarely escaped.

**Fig. 1:**
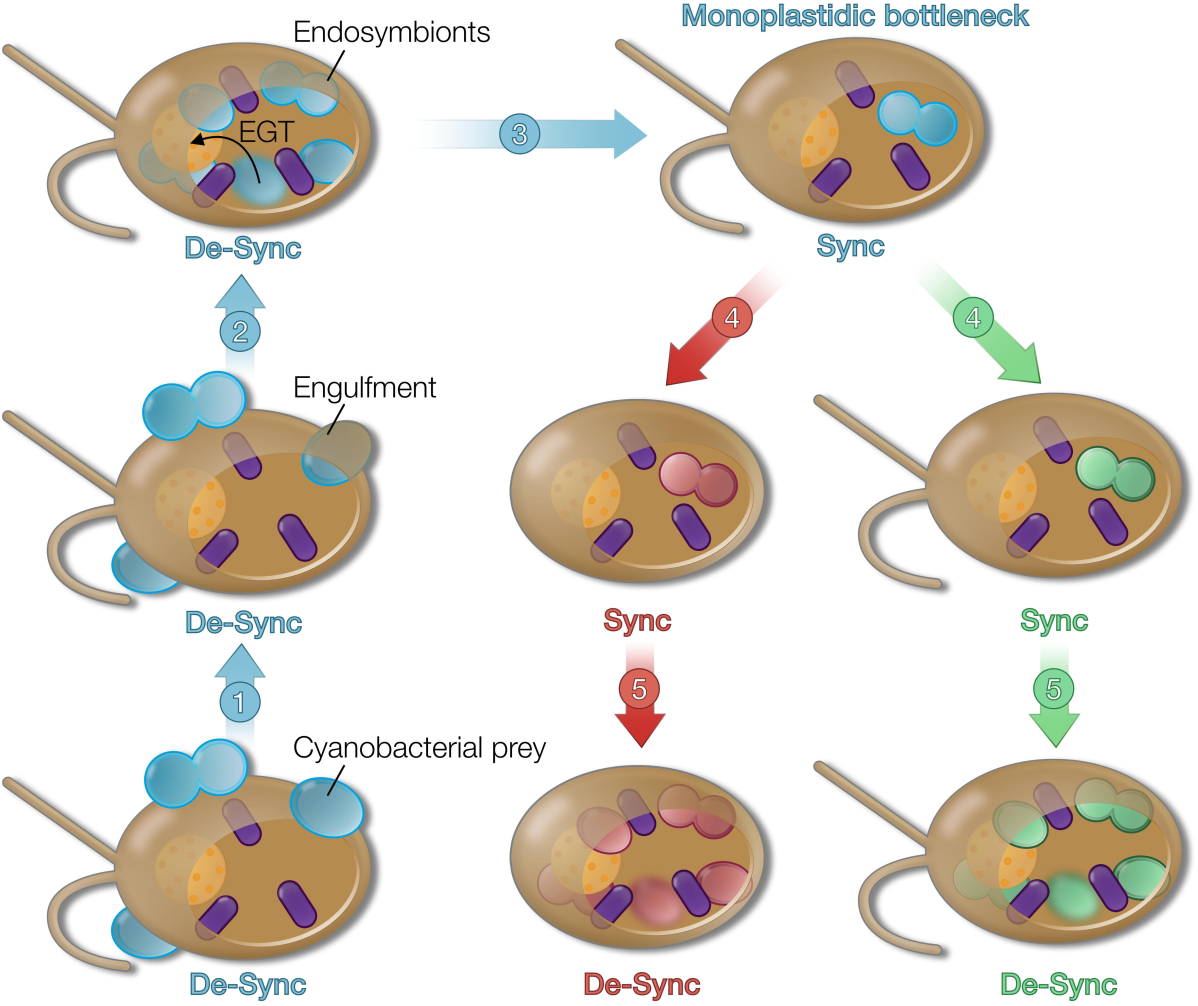
Plastid origin and the monoplastidic bottleneck. Plastids evolved from free-living cyanobacteria (cyan blue) that were originally engulfed by a heterotrophic protist (the first common ancestor to all Archaeplastida with a nucleus in yellow and mitochondria in purple) as prey for the purpose of digestion. After phagocytotic uptake and during the early stages of endosymbiont evolution, the division of the cyanobionts was not yet synchronized (“de-sync”, step 2) with that of the protist host and multiple endosymbionts per nucleus were present. At some point during their co-evolution, the division of the plastid became synchronised (“sync”) with that of the host (step 3), also through what we call the monoplastidic bottleneck, which is immanent in all three lineages (Glaucophyta, Rhodophyta and Chloroplastida) that later evolved. Representatives of the red and green lineage (4) escaped that monoplastidic bottle neck independently (step 5) and the division of plastids in their cytosol no longer depends on the simultaneous division of the nucleus and host cell. This situation is reminiscent of the “De-Sync” situation early on during plastid evolution.

## The monoplastidic bottleneck

A plant or algal cell needs to assure the vertical inheritance of its plastids and mitochondria — the alternative is lethal. There are two possible solutions for controlling the inheritance of at least one organelle (of each type) by the offspring. The first is the presence of numerous organelles in the cytosol resulting in a rather passive inheritance based on a stochastic distribution, which might be the case for mammalian mitochondria (cf. Mishra and Chan, 2014) but always carries the risk of random failure. This can maybe be tolerated in multicellular organisms to a degree, but is likely selected against in single celled eukaryotes. The second solution is a synchronization of organelle and nuclear division and controlled distribution of compartments during cytokinesis. Since multiple endosymbionts were likely key for the endosymbiont to organelle transition to foster EGT events, the first option appears the go-to answer, but only for the earliest stages of plastid evolution. Basal branching algae of the red and green lineage are all monoplastidic (Fig. 2), and a few nonvascular land plants are monoplastidic at least during some stages of their life cycle (Brown and Lemmon, 1990; Vaughn et al. 1992). Maybe all glaucophytes are monoplastidic, too, but here the status quo is more involved and we shall later see why. Worth an extra look are those species that harbour not one or many, but two plastids.

**Fig. 2:**
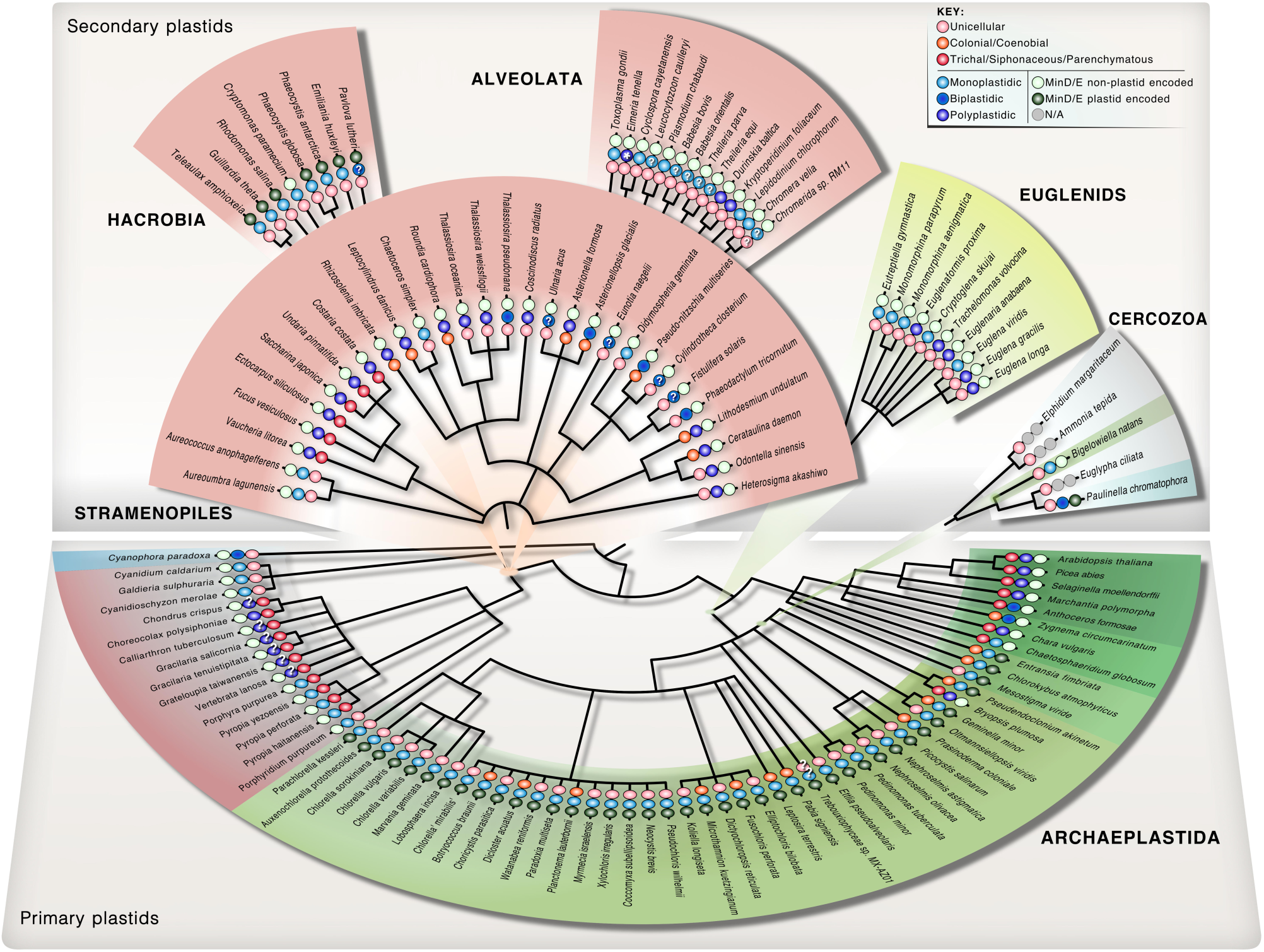
Plastid genomes of polyplastidic algae lack minD and minE. Cladograms of the diversity of photosynthetic eukaryotes based on NCBI Taxonomy. Coloured gradients in the background depict the phylogenetic association of the plastids. The bottom cladogram shows the primary plastid housing Archaeplastida, whose plastids trace back to the monophyletic and endosymbiotic incorporation of a cyanobacterium. Archaeplastida are comprised of the Glaucophyta (blue gradient), Rhodophyta (red) and Chloroplastida (green). Through (serial) secondary endosymbiosis of an early branching rhodophyte, the distantly related Stramenopiles, Hacrobia and Alveolata became secondary red meta-algae (salmon coloured). Secondary green algae arose by two independent events, once acquiring an alga branching basal in all Chlorophyta in case of the Euglenids (chartreuse yellow) and once acquiring a basal-branching ulvophyte in case of the cercozoan Chlorarachniophytes (light green). The Cercozoa also include the only independent primary plastid acquisition, that of the chromatophore, in *Paulinella* (cyan). The entire plastid dataset of NCBI (February 2016; genomic and protein) was screened using a BLASTp and tBLASTn approach and *G. theta* MinD and MinE as queries. Published information on the morphology of the displayed organisms was screened for (i) unicellular, morphologically colonial (coenobial) or “multicellular” growth (including trichal, siphonaceaous, and parenchymatous), and (ii) plastid number (labelling see key; references can be found in Table S1). Not displayed is the plethora of sequenced land plant plastid genomes (five representatives are shown) as well as Chlorophyceae, for which no plastid-encoded MinD or MinE homolog was detected. Note that siphonacous algae are also labelled as multicellular. The asterisk (*) indicates that *Eimeria tenella* forms sporozoites and merozoites that can have more than one apicoplast and highly polyplastidic schizonts (cf. Ferguson et al. 2007).

Among several independent groups, we find representatives with two plastids per nucleus (biplastidy). Biplastidy could be seen as an intermediate stage between housing strictly one or multiple plastids per cell and nucleus, but what, if any, really is the difference between mono- and biplastidic cells and is there a reason for this number? During cell division, a monoplastidic cell must become at least transiently biplastidic to pass on a plastid to its daughter cell. Biplastidy could hence be the result of a simple shift in plastid division from shortly prior to cytokinesis to just after cytokinesis. This seems to have occurred multiple times independently, too. For example, hornworts tend to have one or two plastids per cytosol (therefore labelled biplastidic in Fig. 2; cf. Vaughn et al., 1992) and usually branch at the base of the primarily polyplastidic land plants (Wickett et al., 2014). Furthermore, among the stramenopiles, which contain many polyplastidic species, biplastidy is frequently observed (Fig. 2). Intriguingly, a similar situation is observed in the much younger case of plastid acquisition, namely in *P. chromatophora,* a thecate amoeba that has been co-evolving with is cyanobacterial endosymbiont for about 60 million years (Nowack, 2014). *P. chromatophora* harbours 2 chromatophores, one of which is transmitted to the daughter cell after cytokinesis (Nomura et al., 2014; Nowack, 2014). This mode of inheritance also applies to its relatives such as *P. longichromatophora* (Kim and Park, 2016). The presence of only one chromatophore would drastically diminish the chances of EGTs to occur, trapping *Paulinella* in endosymbiont evolution early on. There might be a lesson to learn here and biplastidy might represent some kind of evolutionary compromise, a compromise between the synchronization of endosymbiont and host also through EGT, and the possibility to lose one organelle due to lysis. Biplastidy is far more prevalent than simple chance would suggest and future research is needed to clarify its evolutionary and molecular significance.

Monoplastidy is the common and ancestral character state of plastid-bearing eukaryotes. It tells us that during the endosymbiont to plastid transition plastid number per cell was reduced to one and probability suggests this ‘monoplastidic bottleneck’ occurred before the split into the three main archaeplastidal lineages (Fig. 1). The archaeplastidal ancestor was also a quite likely a heterotrophic single-celled eukaryote, a protist, and hence evolution selected for monoplastidy to secure proper vertical inheritance of the photosynthetic organelle through the synchronization of host and plastid division. Support comes from the finding that both in *Cyanophora paradoxa* and the red alga *Cyanidioschyzon merolae*, plastid and nucleus division is coordinated in a manner that mitosis only commences after successful plastid division (Sumiya et al., 2016). One can conclude that within the archaeplastidal ancestor a common set of regulatory factors was implemented that orchestrates plastid division and synchronizes it with the host cell cycle. In the following, we inspect what these factors might be and what they mean with regard to the evolution of the plastid division machinery.

## Plastid division and number across archaeplastidal cytosols

Plastids are of cyanobacterial origin (Mereschkowsky, 1905; Cavalier-Smith, 1982; Dagan et al., 2013; Ponce-Toledo et al., 2017; Fig. 1) and hence inherited the backbone of their division (i.e. fission) machinery from their cyanobacterial ancestor (Miyagishima and Kabeya, 2010). Cyanobacterial division differs in some components from that of other bacteria, but is ultimately also based on the physical constriction of the cell carried out by a contractile ring (Miyagishima et al., 2005). Formation of this contractile ring hinges upon the activity of a self-assembling GTPase called <u>F</u>ilamentous <u>t</u>emperature-<u>s</u>ensitive protein <u>Z</u> (FtsZ; Bi and Lutkenhaus, 1991; de Boer et al., 1992; Osawa et al., 2008; de Boer, 2010). FtsZ is the primary component of the division ring that forms in the bacterial cytosol and a whole array of accessory components either controls formation of the Z ring or get recruited after the Z ring is formed (Adams and Errington, 2009). Plastids have inherited many of these components, including FtsZ (Strepp et al., 1998; Miyagishima et al., 2014a; Osteryoung and Pyke 2014), but they now act in concert with the host cell cycle (Sumiya et al., 2016), not least because most of them are now nuclear-encoded (Miyagishima et al., 2012).

During plastid division in Chloroplastida and Rhodophyta, the gene expression profiles of some genes associated with plastid division share the same pattern (Miyagishima et al., 2012). This includes *ftsZ, DRP5B* (<u>d</u>ynamin-<u>r</u>elated <u>p</u>rotein) involved in organelle scission from the outside, and also the *<u>mini</u>cell* gene *minD,* which regulates the positing of the FtsZ-based ring on the stromal side of the inner plastid membrane (Fujiwara et al. 2008; Osteryoung and Pyke 2014). In case of the chloroplastidian *minD* it made no difference to the oscillation pattern, whether it was nuclear or plastid-encoded; although in the latter case *minD* expression seemed light- and not cell cycle-regulated (Miyagishima et al., 2012). Plastid division genes *(ftsZ*, *ftsW* and the <u>sep</u>tum development gene *sepF)* of the glaucophyte *Cyanophora paradoxa* in comparison, however, experience only very little expression fluctuations (Miyagishima et al., 2012). Notwithstanding, *C. paradoxa* and *C. merolae* share the same mechanism that allows mitosis to commence only after plastid division is completed (Sumiya et al., 2016). Yet, the differences in oscillation of division factors might indicate that the plastid division in *Cyanophora* is somewhat distinct from that of the other Archaeplastida. How so?

Glaucophytes are the deepest diverging clade of the Archaeplastida (Burki, 2014). There are only 15 glaucophyte species described (Guiry, 2012), but more likely exist (Jackson et al., 2015; Takahashi et al., 2016). The most basal-branching glaucophyte genus is *Cyanophora* (Chong et al., 2014). *Cyanophora* cells tend to harbour 1 or 2 plastids, and if 2 then they are semi-connected as if frozen in the act of division (Jackson et al., 2015). The other well-characterised glaucophyte genera such as *Cyanoptyche, Gloeochaete* or *Glaucocystis* are polyplastidic (Jackson et al., 2015), but *Glaucocystis*’ plastid morphology warrants attention. *Glaucocystis* has two stellate cyanelle complexes (Schnepf et al., 1966). These cyanelle complexes consist of single cyanelles that, however, are kept together through unknown means in a pack of two (Schnepf et al., 1966). Either way, the basal position of *Cyanophora* in a clade of polyplastidic algae further supports that housing two (connected) plastids per cytosol might be an intermediate stage between mono- and polyplastidy. *Glaucocystis*’ bundling of plastids might be a relic of this transition. Glaucophytes are generally considered to have retained ancestral features of the earliest Archaeplastida (Fathinejad et al., 2008; Steiner and Löffelhardt, 2011; Facchinelli et al., 2013), including a thick peptidoglycan layer between the two membranes of the organelle (Steiner et al., 2001).

## Plastid division and the ancestral peptidoglycan layer

The cyanobacterial progenitor of the plastid had a layer of peptidoglycan (PG), separating its inner and outer membrane. This is evident by the retention of the PG (or murein) layer by the plastids (cyanelles) of glaucophytes (Steiner et al. 2001) and the recent identification of a thin PG layer with D-amino acids surrounding the plastids of *Physcomitrella* (Hirano et al., 2016). Until the latter discovery, there was a consensus that this murein layer had been lost early during plastid evolution after the green and red lineage had diverged from the glaucophytes, also because the PG layer is thought to interfere with protein import of nuclear-encoded plastid proteins (Steiner and Löffelhardt, 2002). Nonetheless, the presence of PG-synthesizing enzymes in photosynthetic eukaryotes (Machida et al., 2006; Garcia et al., 2008; Takano and Takechi, 2010) was always suspicious and the PG-layer known to be somehow associated with plastid division. Could the retention of even only a thin murein layer be associated with some kind of regulation of plastid division?

Independent lines of evidence connect the PG-layer with the regulation of plastid division. PG-inhibiting antibiotics such as ampicillin or fosfomycin effect plastid division and morphology in streptophyte algae (Matsumoto et al., 2012), lycophytes (Izumi et al., 2008) and mosses (Katayama et al., 2003). Plastid division was also altered in *Physcomitrella* lines, whose PG synthesizing enzymes were knocked-out (Homi et al., 2009). Such effects are, however, not observed in any euphyllophyte (Takano and Takechi, 2010) and might thus be part of the specific evolutionary changes the plastids of higher vascular plants, the embryoplast *(cf.* de Vries et al., 2016), experienced. Even a link between the complete loss of the PG layer and the switch from mono- to polyplastidy in embryophytes is feasible, but alone cannot be the reason considering the absence of a PG layer in rhodophytes that however are mostly monoplastidic (Fig. 2). In bacteria, loss of the PG layer and loss of FtsZ correlate (Miyagishima et al., 2014a), which makes sense considering that the guidance of PG layer synthesis is among the main functions of FtsZ (de Pedro et al., 1997; Aaron et al., 2007; Typas et al., 2012) that together can influence bacterial morphology (Kysela et al. 2016). Yet, plastid division still involves FtsZ in all Archaeplastida analysed thus far (Yang et al., 2008; Miyagishima et al., 2012). FtsZ is, however, not essential for plastid division (Schmitz et al., 2009; Miyagishima et al. 2014a). Plastids of *ftsZ* knockout (KO) lines — including double and triple knockouts — still divide, but intriguingly their cells harbour a reduced number of plastids per cell (Schmitz et al., 2009). This suggests that FtsZ is of major importance regarding fine-tuning, but not essential for the functionality of plastid division.

## Amenability by external forces and the ‘inside first’ in plastid division

Plastid division starts within the organelle (Miyagishima et al., 2014a), the reason for which lies with the plastids’ cyanobacterial origin. The cyanobacterial division machinery works from the cytosolic face of the plasma membrane first (Errington et al., 2003), which makes sense considering the alternative would be the secretion of a division machinery into the periplasm and environment to act on the exterior side of the two membranes. But with the cyanobacterial ancestor resting in the cytosol of the host cell, and the concomitant reduction or even loss of the PG layer, plastids became accessible to and amendable by external forces. The most palpable of these external forces is the outer division ring formed mainly by a protein of the dynamin family (cf. Yang et al., 2008; Miyagishima and Kabeya, 2010) and the latter’s origin and function is connected to the origin of mitochondria (Purkanti and Thattai, 2015; Gould et al., 2016).

Dynamins are eukaryote-wide conserved GTPases that exercise mechanical forces and some dynamins mediate the contraction and scission process during organelle division (McFadden and Ralph, 2003; Purkanti and Thattai, 2015; Leger et al. 2015). Dynamin-mediated plastid division arose in the common ancestor of Rhodophyta and Chloroplastida, as it is present in all studied members of the red and green lineage, but not glaucophytes (Miyagishima et al. 2014b). Interestingly, carrying mutations in genes coding for inner division machinery components, such as *ACCUMULATION AND REPLICATION OF CHLOROPLASTS 6 (ARC6;* a J-domain protein homolog to the cyanobacterial Ftn2 [cf. Vitha et al., 2003]), has a more pronounced influence on plastid number (Fig. 3) than mutations in components acting on the outside (cf. Robertson et al., 1996; Sakaguchi et al., 2011). Moreover, proplastid division is not affected in dynamin knockout lines (Robertson et al., 1996). Yet, regardless of whether outer or inner division components are manipulated, proplastid division still occurs (cf. Miyagishima et al., 2014a). Knockout of components reduces plastid number (Fig. 3), in land plants even to the degree that it only allows proplastid division (Miyagishima et al., 2014a). All things considered, plastid division is overall quite robust and that these few plastids can still divide pays tribute to the framework established at the monoplastidic bottleneck. Controlled polyplastidy arose by complementing this robust monoplastidic framework with additional layers of regulation. Dynamin hence represents one of these additional layers of control the host exercises over organelle division and which was originally implemented to work on the mitochondrion.

**Fig. 3:**
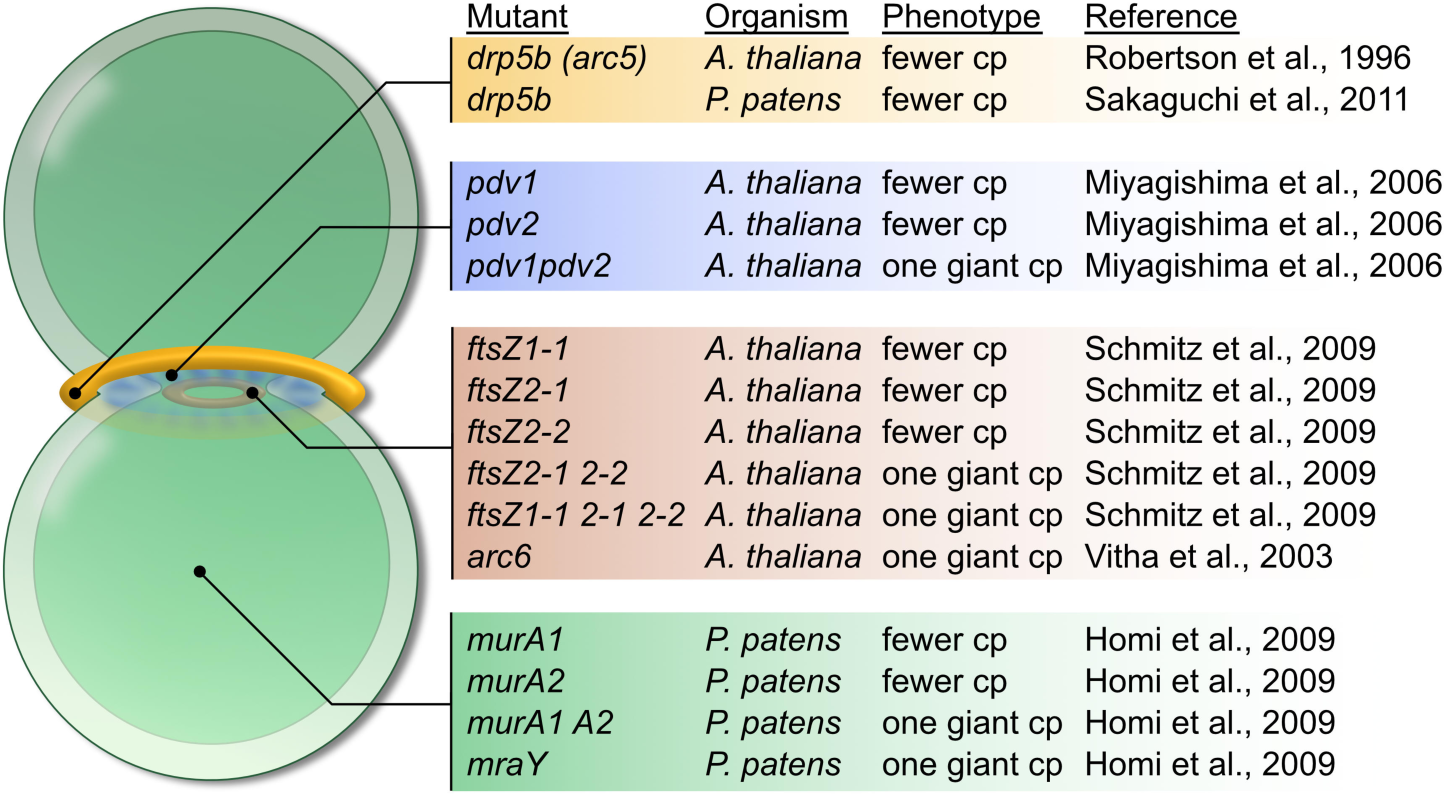
Single mutations can cause strong alterations in plastid number per cell. Schematic drawing of a dividing plastid, depicting outer division components (yellow), the land plant-specific PDV proteins (blue), and the inner division components (red). On the right, the respective division components and published information on the phenotypes of plastid-division mutants are listed (in the same colour-code as on the left).

Several times independently, components of the plastid division machinery were lost or modified (Miyagishima et al., 2014a). So far, we have discussed a handful of genes and properties that might have been crucial for the orchestration of plastid division. This raises the question of whether maybe just a few key factors determine whether one or many essential organelles are present in a cell. Maybe, but it is complicated. Some studies have shown that the knock-out of single genes can transform a polyplastidic embryophyte cell to one carrying a single embryoplast (Fig. 3). These monoplastidic mutants often harbour a single plastid massively increased in size, reminiscent of large algal plastids such as that of *Chlamydomonas,* whose plastids take up about half of the cell volume (cf. Gaffal et al., 1995). This response in increased compartment size is likely communicated through shuttling of the REC (<u>r</u>educed <u>c</u>hloroplast <u>c</u>overage) proteins between the cytosol and the nucleus (Larkin et al., 2016). Such large single plastids are observed when plastid division is impaired, for example in *arc6* (Vitha et al., 2003) and *ftsZ* (Schmitz et al., 2009) knockouts, and even in PG synthesis *murA* and *mraY* knockout lines of *Physcomitrella patens* (Homi et al., 2009; Fig. 3). What this means is that it takes only a single gene to revert polyplastidy back to (macroplastidic) monoplastidy.

## *minD* and *minE:* markers for the evolution of complex plastids

Plastid division requires the interplay of two genetic compartments, that of the plastid and the nucleus. While in land plants all plastid division proteins are nuclear-encoded, the situation is quite different in algae: their plastid genomes can include the genes for *ftsI, ftsW, sepF, minD* and *minE* (Miyagishima et al., 2012). Recently, we asked the question of whether the absence of *minD* and *minE* from the plastid genome could be a prerequisite for the evolution of polyplastidy (de Vries et al., 2016). We screened all 999 species of Archaeplastida and secondary plastid-bearing lineages for which plastid genomes were available at NCBI (as of February 2016) for (i) the presence of *minD* and *minE* genes in their plastid genomes and (after excluding most of the numerous chlorophyceaen and land plants species with sequenced plastid genomes) the morphological characters of 131 species for (ii) mono-, bi- or polyplastidy, and (iii) unicellularity, colonial morphology or multicellularity (Fig. 2). Algae that encode genes for *minD* and *minE* in their plastid genome are exclusively monoplastidic and limited to primary green Chlorophyta (which is also absent in the secondary green chloroplast genomes of Euglenozoa that are often polyplastidic) and secondary red Hacrobia (Fig. 2). The latter is peculiar.

Only plastid genomes of Hacrobia (haptophytes+cryptophytes; cf. Okamoto et al., 2009, but see also Burki et al. 2012) and *P. chromatophora* encode *minD* and *minE.* In fact, among all Archaeplastida, cryptophyte plastid genomes are the only ones to still encode *minE.* The deep branching of cryptophytes among photosynthetic eukaryotes (Stiller et al., 2014; Burki, 2014) and certain trades associated with protein import (Gould et al., 2015) suggest they represent an ancestral state, which the coding of *minD* and *minE* by the plastid genome further supports. It was recently suggested that haptophytes acquired their plastids from ochrophytes (stramenopiles) through quaternary endosymbiosis (Stiller et al., 2014), but in light of *minD/E* distribution this is cause for reflection. No stramenopile plastid genome encodes either a *minD* or *minE* homolog (Fig. 2), so that haptophytes would have acquired a stramenopile with a plastid that was unlike any of those known today. Not impossible, but considering in addition that the nuclear-encoded *minD* and *minE* of stramenopiles are likely of mitochondrial origin (Leger et al., 2015), speaks against a quaternary endosymbiotic origin of the haptophyte plastid.

If none of the primary red algae has a *minD* or *minE* homolog found in its plastid genome, where does the one found in the secondary red Hacrobia come from considering that EGT is usually a one-way-ticket (cf. Martin et al., 1998). The Cyanidiales are on the deepest branch in red algal phylogenies (Yoon et al., 2006). Among the Cyanidiales, *Galdieria sulphuraria* is the only red algae that still harbours some remnants of a *minD* homolog *(cf.* Leger et al., 2015; see Fig 2). *G. sulphuraria* diverged very early from the other Cyanidiales (Yoon et al., 2006), which means that secondary red plastids must have been acquired very early on during rhodophyte evolution, likely from an extinct or non-sampled red algal lineage, and before these genes were lost or transferred to the host nucleus. Plastid-encoded *minD* and *minE* hence offer additional good markers for tracing the complex evolutionary trajectory of red meta-algae.

## Multicellularity and polyplastidy are not coupled

Plastids evolved in a unicellular eukaryote. Indeed, in almost any lineage, basal branching algae are unicelluar (Fig. 2). The recent placement of the multicellular (i.e. a mass of unicells in a gel matrix) Palmophyllaceae at the base of all Chlorophyta (cf. Leliaert et al., 2016) does not change this observation, as palmophyllaceaen multicellularity likely represents a derived character state. If one paints a simplistic trajectory of algae and plant evolution, one might get the impression that there was a clear overall increase in morphological complexity and that polyplastidy evolved as a by-product. This is not the case, as the green and red lineage evolved multicellularity multiple times independently (Lewis and McCourt, 2004; Leliaert et al., 2012; Cock and Collén, 2015). Ancestral character state inferences even suggest that multicellularity arose in just single algal classes several times independently, as observed for example within the ulvophytes (Cocquyt et al., 2010). However, multicellularity and polyplastidy do not go hand in hand. Based on the characters of basal-branching streptophytes (*Mesostigma viride;* [Marin and Melkonian, 1999]), and chlorophytes (the prasinophytes; Leliaert et al., 2012), polyplastidy was not a feature of the common ancestor of all Chloroplastida (cf. Leliaert et al., 2011; Leliaert et al., 2012).

Polyplastidic, multicellular species are not limited to streptophytes. Non-streptophyte examples include rhodophytes such as *Gracilaria* or *Choreocolax* (Callow et al., 1979; cf. Schmidt et al. 2010) or phaeophytes such as *Ectocarpus* (cf. Charrier et al., 2008, Fig. 2). Monoplastidic, multicellular algae can be found across a broad taxonomic range, too, including the chlorophytic Palmophyllaceae (cf. Zechman et al., 2010; cf. Lelieart et al., 2016) or the rhodophytes *Porphyra* and *Pyropia* (Sutherland et al., 2011). Polyplastidic, unicellular algae seem to mainly occur among secondary-plastid housing organisms such as the euglenophyte *Euglenaformis* [cf. Bennett et al., 2014] or the stramenopile *Heterosigma* (Hara and Chihara, 1987). A curious type of polyplastidy occurs in coenocytic (siphonaceous) algae such as *Vaucheria*. Belonging to the heterokontophytes (whose complex plastids are surrounded by four membranes, the outermost thought to be continuous with the endoplasmic reticulum [Gould et al. 2015]), one would suspect that their plastids occur in a complex with the nucleus, each complex containing one nucleus and two plastid lobes (cf. Apt et al., 2002). Could it be that they are effectively biplastidic, albeit a single *Vaucheria* cell can contain thousands of nuclei-plastid complexes (Ott 1992)? We are not aware of any in-depth analysis on this subject and apart from a RNAseq analyses that revealed a spatial mRNA distribution in the siphonaceous ulvophyte *Caulerpa taxifolia* (Ranjan et al., 2015), little molecular work is carried out on siphonaceous algae in general. Studying nucleus-plastid communication in a coenocytic system using modern tools might therefore hold some surprises.

## Lessons from mitochondrial inheritance

Mitochondria and plastids face similar challenges. The remaining organellar genomes are first of all in danger of mutational meltdown due to Muller’s ratchet (Lynch et al., 1993; Lynch, 1996; Martin and Herrmann, 1998), concomitant with an elevated risk of accumulating mutations induced by reactive oxygen species (Aro et al., 1993; Apel and Hirt, 2004; Balaban et al., 2005). One can assume that evolution has favoured those organisms, which evolved scrupulous mechanisms that protect the integrity of the rudimentary organellar genomes. For mitochondria, there are many ideas and studies on how this is implemented *in vivo,* not least because inheriting a mitochondrion with a deleterious mutation can cause human diseases such as Leber’s hereditary optic neuropathy or MERRF syndrome (Holt et al., 1989, Larsson et al., 1992) and the mutational baggage accumulating over a lifetime in mitochondrial DNA (mtDNA) has repeatedly been associated with aging (Harman, 1956; Harman, 1972; Balaban et al., 2005).

Assuring that mutated mitochondria do not spread in animal populations could be achieved by (i) preventing those organelles that are subject to inheritance from accumulating possibly deleterious mutations, i.e. keeping organelles in mint condition, (ii) assuring that only mint condition organelles are inherited, or (iii) a combination of the two. There is some compelling evidence for the former: de Paula and colleagues (2013) found that mitochondria in the ovaries but not sperm cells of fruit flies and zebrafish stay bioenergetically inactive (i.e. no ATP-synthesis through an electron transport chain), hereby significantly reducing the formation of DNA-damaging ROS. It is thought that these inactive mitochondria then serve as templates for the next generation. Regarding the second option, there exists a concept of a mitochondrial genetic bottleneck occurring in animal germ cells. This genetic bottleneck is achieved by reducing the intracellular population of mitochondria/mtDNAs, where the mutational load in this reduced set of mtDNAs is thought to bear faster and more pronounced on the physiology of a given population of germ cells, inducing a purifying selection that favours germ cells with fit mitochondria (Bergstrom and Pritchard, 1998). Although various sorts of experimental data have been gathered to refine the genetic bottleneck model, there seems to be no consensus on when and how this bottleneck occurs. Independent studies have found a significant reduction in mtDNA at some point during germline development in mice (Cree et al., 2008; Wai et al., 2008), although these data have been challenged (Cao et al., 2009). A recent study found that selective proliferation favoured non-mutated (i.e. “wildtype“) mtDNA in *Drosophila* and endorsed selection at the organellar level instead of at a cellular level (Hill et al., 2014). Regardless of the details of these controversies, a genetic bottleneck in mitochondrial inheritance likely exist (Stewart and Larsson, 2014).

There are obviously limits in transferring this animal-centric information onto the plastids of plants and algae. First, in many plastid-housing eukaryotes organellar mutation rates are quite low (Smith, 2015), which could, however, also be a result of the mechanisms outlined here. Second, the lack of immediate (i.e. embryonic) separation of germ and soma cells. Third, most algae are unicellular. The formulation of models on such a separation are therefore tricky, although a mechanism that ensures the inheritance of only fit organelles in protists is generally feasible. In budding yeast, especially those mitochondria with a high Δψ (indicative of bioenergetic vigour) are inherited because they localise to the budding site (Higuchi-Sanabria et al., 2016). Furthermore, the genetic mechanisms discussed also work in unicellular and monoplastidic — but nonetheless multiple plastid genome copies-bearing — species such as *Chlamydomonas* (VanWinkle-Swift, 1980; Birky, 2001).

The number of proplastids in a seed is rather small (Possingham, 1980) and that number might be actively reducedMogensen 1996). Could there be a connection between the (ancestral) monoplastidic bottleneck and the genetic bottleneck during organelle inheritance, especially in polyplastidic organisms? One cannot really tell, but there are numerous mechanisms to selectively restrict plastid numbers in germ cells and plants are generally very efficient at obtaining homoplasmy after several rounds of cell divisions (Greiner et al., 2015). Some basal branching embryophytes were described to be monoplastidic in those cells that are relevant for spore production (Brown and Lemmon, 1990; see also Smith et al. 2011) or even in their meristematic tissue in case of some lycophytes (Brown and Lemmon 1984; 1985). Spores (and gametes;Maier et al. 1997) of the multicellular and polyplastidic brown alga *Ectocarpus siliculosus* are monoplastidic, too (Baker and Evans, 1973), speaking for a potentially analogous need for monoplastidy during *Ectocarpus*’ reproductive cycles. Quality control mechanisms established at the evolutionary-early time of the monoplastidic bottleneck — that evidently happened — would benefit a genetic bottleneck during plastid inheritance. The reduction of plastid number in gametes and spores to one we conclude, is a required reversion back to monoplastidy. To what degree some of these functions are conserved and whether they still act in seed plants remains to be determined, but surely presents a worthwhile endeavour.

## Conclusion

Most might conceive the presence of dozens or hundreds of plastids per cell the norm, but some non-vascular plants and especially algae tell us otherwise. There exists a peculiar monoplastidic bottleneck in algae and plant evolution, whose reason and extend yet remains to be fully explored. The monoplastidic bottleneck occurred in the ancestor of Archaeplastida and at which time the majority of the regulatory mechanisms that govern plastid division (and control plastid number) were implemented. Most of the latter rest upon the modification of the fission machinery the plastid brought along, which were later complemented by the host through proteins such as those of the dynamin family and the <u>P</u>LASTID <u>D</u>IVISION <u>P</u>ROTEINS with the emergence of land plants. To a degree plastid inheritance follows principles that also apply to mitochondria (rarely studied simultaneously) and both are guided by uniparental inheritance that is, however, far more involved regarding the organelle of cyanobacterial origin. Whether uniparental inheritance of plastids is associated with the monoplastidic bottleneck is hard to say, because a systematic study on the topic remains to be conducted. From what we can tell, the presence of a monoplastidic checkpoint even in higher embryophytes (maybe during zygote formation) cannot entirely be ruled out. If it no longer exists in embryophytes, it begs the question of how they managed to escape monoplastidy and whether a similar or a radical different solution evolved in those red algae that escaped the monoplastidic bottleneck independently. The occurrence of polyplastidy coincides with a complex and macroscopic morphology, both in rhodophytes and streptophytes, underscoring the impact of this transition. A testable prediction from our rationale is that the molecular regulation of plastid division in monoplastidic plant cells is homologous to that established at the monoplastidic bottleneck event. Comparative studies hold the potential to uncover the molecular chassis that was established during the monoplastidic bottleneck and to what degree it continues to influence and define plastid inheritance in land plants.

## Acknowledgements

JdV (VR 132/1-1) and SBG (GO1825/4-1 and CRC1208) are grateful for the financial support provided by the German Research Foundation. We thank Klaus V. Kowallik for many useful comments, as well as Pavel škaloud (Charles University, Czech Republic), Monique Turmel (Université Laval, Canada), Eric W. Linton (Central Michigan University, USA), Matthew S. Bennett (Michigan State University, USA) and Robin Matthews (Huxley College of the Environment, USA) for the kind and informative correspondences.

## Author contribution

JdV and SBG together conceived and wrote the manuscript.

